# Pattern similarity and connectivity of hippocampal-neocortical regions support empathy for pain

**DOI:** 10.1101/811935

**Authors:** Isabella C. Wagner, Markus Rütgen, Claus Lamm

**Author notes:** Correspondence: Isabella C. Wagner, Social, Cognitive and Affective Neuroscience Unit, Department of Basic Psychological Research and Research Methods, Faculty of Psychology, University of Vienna, Liebiggasse 5, 1010 Vienna, Austria.

## Abstract

Empathy is thought to engage mental simulation, which in turn is known to rely on hippocampal-neocortical processing. Here, we tested how hippocampal-neocortical pattern similarity and connectivity contributed to pain empathy. Using this approach, we analyzed a data set of 102 human participants who underwent functional MRI while painful and non-painful electrical stimulation was delivered to themselves or to a confederate. As hypothesized, results revealed increased pattern similarity between fist-hand pain and pain empathy (compared to non-painful control conditions) within the hippocampus, retrosplenial cortex, the temporo-parietal junction and anterior insula. While representations in these regions were unaffected by confederate similarity, pattern similarity in the dorsal MPFC was increased the more dissimilar the other individual was perceived. Moreover, hippocampal connectivity with regions engaged in first-hand pain was also increased during pain empathy, during which hippocampal coupling with the fusiform gyrus positively scaled with self-report measures of individual perspective taking skills. These findings highlight that shared representations and interactions within a hippocampal-neocortical network support pain empathy. This potentially reflects memory-based mental simulation processes, which seem partially modulated by personality traits and the perceived similarity of the other individual in pain.

---

Empathy describes the ability to share the emotional state of another person and is crucial for successful social interactions. The so-called “shared representations account” suggests that empathy for an affective state engages similar neuronal processes as experiencing the affective state directly (Zaki et al., 2016; Lamm et al., 2019). In line with this assumption, previous studies implicated the dorsal anterior cingulate cortex (also termed anterior mid-cingulate cortex; dACC/aMCC) and the anterior insula in the first-hand experience of pain and in pain empathy (Fan et al., 2011; Lamm et al., 2011; Rütgen et al., 2015b, 2015a, 2018; Marsh, 2018). Furthermore, a recent animal study revealed neurons within the rat ACC that coded not only for first-hand pain but also fired when rats witnessed a conspecific receiving footshocks (Carrillo et al., 2019). This data, however, stands in contrast to divergent results from multivariate analyses that reported both shared and distinct representations during experienced emotions and empathy (Corradi-Dell’Acqua et al., 2011, 2016; Krishnan et al., 2016), altogether fueling a long-standing debate on the neural underpinnings of empathy (Zaki et al., 2016; Lamm et al., 2019).

Besides affect sharing, empathy is thought to depend on multiple processes, including self-other distinction and mentalizing, i.e., mentally simulating the stance of another person (see Lamm et al., 2019 for a review). Social cognition theories of mental simulation posit that individuals use their own mental states as models to understand the mental states and actions of others (Gallese and Goldman, 1998). This has been associated with neural processing in a set of regions (for a review, see Frith and Frith, 1999, 2006; Uddin et al., 2007; Mar, 2011), focused predominantly on the medial prefrontal and posterior cingulate cortex (MPFC and PCC; Gallagher et al., 2000, 2002; Amodio and Frith, 2006; Saxe and Powell, 2006; Spreng and Grady, 2010), and the temporo-parietal junction (TPJ; Saxe and Kanwisher, 2003; Saxe and Wexler, 2005). The hippocampus and adjacent medial temporal lobe (MTL) structures have well-documented roles in recalling past (Scoville and Milner, 1957; Zola-Morgan and Squire, 1990; Vargha-Khadem, 1997; Frankland and Bontempi, 2005; Rugg and Vilberg, 2012; Kim, 2016) and simulating (future) events (Klein et al., 2002; Buckner and Carroll, 2007; Hassabis et al., 2007a, 2007b; Schacter and Addis, 2007; Hassabis and Maguire, 2009; Mullally et al., 2012). Surprisingly, the potential contributions of these areas to empathic processing have been largely neglected. Initial evidence that links empathy with hippocampal volume stems from studies with patients suffering from traumatic brain injury (Rushby et al., 2016), hippocampal amnesia (Beadle et al., 2013), as well as from a recent developmental study (Stern et al., 2019). Here, we capitalized on hippocampal processing within a sizable sample of healthy participants that engaged in an empathy for pain task.

To pursue this novel focus on hippocampal-neocortical processing, we chose an analysis approach complementary to previous analyses that focused on affect sharing during pain empathy (Rütgen et al., 2015b). One-hundred-and-two participants underwent functional MRI while completing a pain empathy task where painful and non-painful electrical stimulation was delivered to the participant or to a confederate (**Figure 1AB**). Importantly, this task was designed to elicit mental simulation processes while attenuating contributions of mirror neuron activity or motor mimicry. Participants were thus presented with a cue that indicated the intensity of the upcoming electrical stimulation rather than with pictures of the confederate in painful or non-painful situations. First, we hypothesized that if mental simulation contributes to pain empathy, participants should base the evaluation of another individual’s pain on representations of their previous, first-hand pain experiences. This should become apparent through similar neural representations between first-hand pain and pain empathy within the hippocampus and distributed regions important for mental simulation and pain empathy, including the MPFC, PCC, TPJ, dACC/aMCC, and the anterior insula. We quantified the underlying neural representations by deriving the multivoxel pattern similarity across single trials, embedded within a whole-brain representational similarity analysis (RSA) framework. Second, if participants use first-hand pain experiences to evaluate the pain of others, we expected this to be dovetailed with the recall of recent information from memory. To this end, we tested increases in task-based connectivity during pain empathy between the hippocampus and neocortical regions that should also be involved when experiencing pain first-hand (Frankland and Bontempi, 2005; Bastiaansen et al., 2009). Third, because mental simulation was shown to depend on perceived similarity with the other individual (Mitchell et al., 2005, 2006; Tamir and Mitchell, 2010, 2013; Majdandžić et al., 2016; Thornton et al., 2019), we further took into account individual ratings of confederate similarity and explored its relationship with neural pattern similarity. Lastly, we stratified our results with trait measures of empathic abilities. By means of this integrative approach, we expect to shed new light on the engagement of hippocampal-neocortical regions, and thus the role of memory-based mental simulation processes in pain empathy.

**Figure 1.**
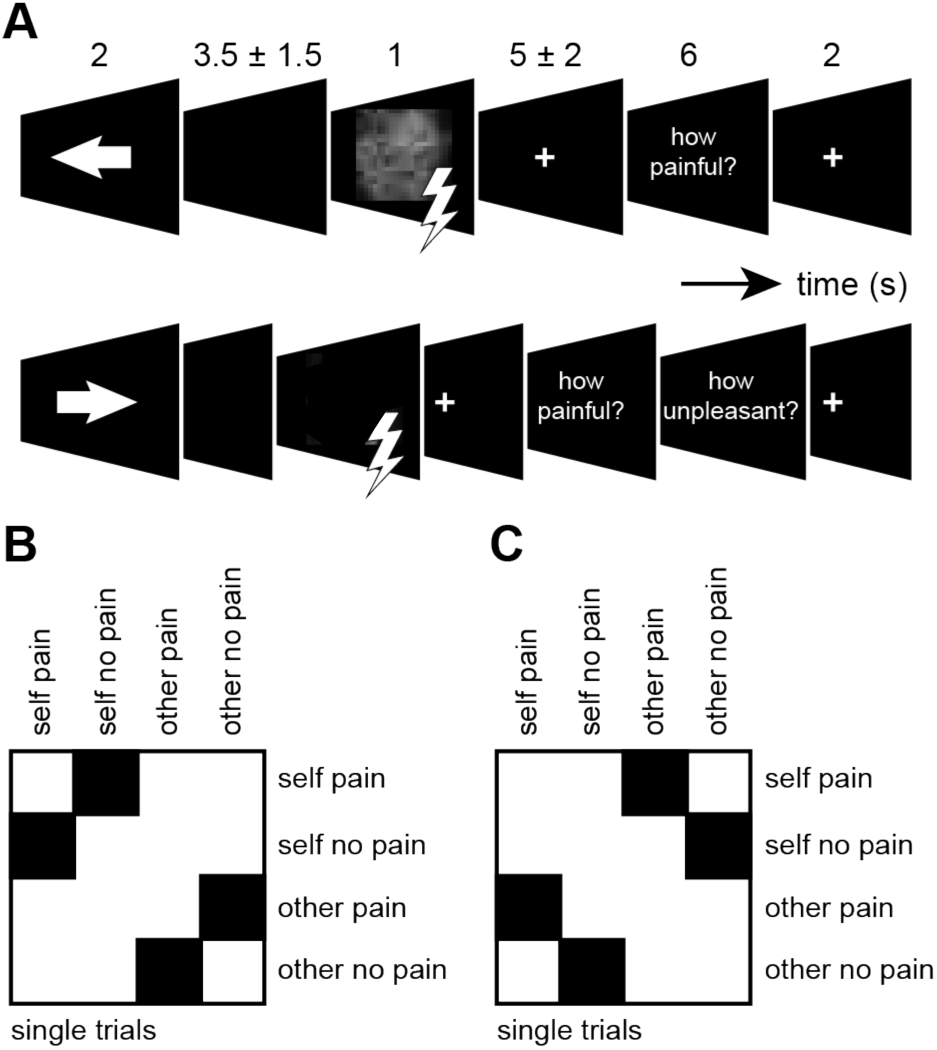
Pain empathy task and representational similarity analysis (RSA). **(A)** Examples for self- and other-directed trials during the pain empathy task. Participants first received a cue if the electrical shock was directed at themselves (upper row) or at the confederate (lower row). Arrow color provided information about the intensity of the upcoming electrical stimulation (red, painful; green, non-painful; not shown here). Participants then saw either a scrambled photo of themselves or a photo of the confederate showing a neutral or painful facial expression during non-painful and painful trials, respectively (note that photographs are not shown in the *bioRxiv* version of this article). After this, participants were asked to rate pain and unpleasantness during one-third of the trials. RSA was performed across single trials of the pain empathy task after trials were sorted according to task conditions (i.e., self pain, self no pain, other pain, other no pain): **(B)** pattern similarity was computed between [self pain × self no pain] and [other pain × other no pain] trials, **(C)** and between [self pain × other pain] and [self no pain × other no pain] trials. Pattern similarity values were extracted from the respective quadrants (marked in black).

## Materials and methods

This study was part of a larger project investigating the effects of placebo analgesia on pain and pain empathy (Rütgen et al., 2015b). In brief, participants had been randomly assigned to a placebo or control group and completed a pain empathy task, as well as an affective touch task (not discussed here), followed by the structural scan and a resting-state period (not discussed here), all inside the MR scanner. We previously reported results from univariate activation analysis of the pain empathy task, comparing placebo and control groups (Rütgen et al., 2015b). Here, we provide a novel analysis focused on pattern similarity and hippocampal connectivity during pain empathy. Separate analyses yielded highly similar results in both subgroups and no significant differences between the groups (not reported further). Since we were interested in generalized contributions of shared representations and mental simulation processes to pain empathy, we thus collapsed our analyses across the two groups.

### Participants

One-hundred-and-two participants were included in this analysis (see Rütgen et al., 2015b for details) (70 females, age range = 19-38 years, mean age = 25). All were right-handed, healthy, had normal or corrected-to-normal vision, and gave written informed consent prior to participation. The study was reviewed and approved by the ethics committee of the Medical University of Vienna (Vienna, Austria).

### Task and procedures

#### Electrical stimulation and pain calibration

Individual intensity values (mA) for electrical stimulation were determined during pain calibration. This involved a staircase procedure where participants were asked to rate pain intensity after every electrical shock (500 ms) using a 7-point scale (1, “perceptible, but no painful sensation”; 7, “extremely painful”). The same scale was used for pain intensity ratings throughout the study. Electrical stimulation was delivered using a Digitimer DS5 Isolated Bipolar Constant Current Stimulator (Digitimer Ltd, Clinical & Biomedical Research Instruments) and a bipolar concentric surface electrode attached to the dorsum of the left hand. Shock delivery was controlled manually using Cogent (version 1.32, www.vislab.ucl.ac.uk/cogent.php).

#### Pain empathy task

During the pain empathy task, participants received a cue (2 s) if the electrical shock was directed at themselves (arrow pointing left, self-directed trial; **Figure 1A**) or at another participant (arrow pointing right, other-directed trial; **Figure 1B**). Additionally, the color of the arrow informed the participant about the upcoming stimulation intensity (red, painful; green, non-painful). The other participant was a member of the experimental team and actually never received any shocks. After a brief delay jittered between 2 and 5 s (mean = 3.5 s), participants saw a photo of the shock recipient on the screen (1 s; self-directed trial: scrambled photo of themselves; other-directed trial: photo of the confederate with painful/neutral facial expression), and a brief electrical shock (500 ms) was delivered (during self-directed trials only). This was followed by a fixation period ranging from 3 to 7 s (mean = 5 s) and affect ratings (6 s) which were collected during one-third of the trials (self-directed pain ratings: “How painful was this stimulus for you”, other-directed affect ratings: “How painful was this stimulus for the other person”, and “How unpleasant did it feel when the other person was stimulated”). Trials were separated with a short fixation period (2 s). In total, participants completed 15 trials per condition (i.e., self-directed painful, self-directed nonpainful, other-directed painful, other-directed non-painful). The task was programmed and presented with Cogent (version 1.32, www.vislab.ucl.ac.uk/cogent.php) and lasted for approx. 16 min.

Stimulation intensities during self-directed trials were set to individually calibrated stimulation intensities related to pain ratings of 1 (i.e., non-painful trial) and 7 (i.e., painful trial) throughout the task. The average stimulation intensities were 0.15 ± 0.14 mA (mean ± SEM; pain intensity rating of 1) and 0.74 ± 0.6 mA (pain intensity rating of 7) during non-painful and painful trials, respectively.

#### Post-experimental ratings and questionnaire data

After MR scanning, participants rated how similar they perceived the confederate, how much they liked the other person, perceived affiliation with the other person, attributed strength, neediness, and agreeableness. Here, we focused on perceived confederate similarity only (“How similar was the other person to you”; 1, “dissimilar”; 9, “very similar”). This rating was not available for one participant and we thus excluded this person from all analyses that involved confederate similarity (i.e., *N* = 101). To assess trait empathy, participants completed the Interpersonal Reactivity Index online prior to the start of the experiment (IRI; subscales personal distress, perspective taking, empathic concern, fantasy; Davis, 1983).

### MRI data acquisition

Imaging data were acquired using a 3 Tesla MRI scanner (Tim Trio, Siemens, Erlangen, Germany) equipped with a 32-channel head coil. We obtained approx. 500 T_2_*-weighted BOLD images during the pain empathy task, using a multiband-accelerated echoplanar imaging (EPI) sequence. Parameters were as follows: TR = 1800 ms, TE = 33 ms, flip angle = 60°, interleaved slice acquisition, 54 axial slices, FOV = 192 × 192 × 108 mm, matrix size = 128 × 128, voxel size = 1.5 × 1.5 × 2 mm. Structural scans were acquired using a magnetization-prepared rapid gradient echo (MP-RAGE) sequence with the following parameters: TR = 2300 ms, TE = 4.21 ms, 160 sagittal slices, FOV = 256 × 256 mm, voxel size = 1 × 1 × 1.1 mm.

### MRI data preprocessing

A detailed description of data preprocessing was reported previously (Rütgen et al., 2015b). In brief, data was processed using SPM12 (http://www.fil.ion.ucl.ac.uk/spm/) and Matlab (The Mathworks, Natick, MA, USA), including slice time correction, spatial realignment, normalization to the Montreal Neurological Institute (MNI) standard space, and spatial smoothing with a Gaussian kernel (6 mm full-width at half maximum, FWHM).

### fMRI data modeling

First, we used Representational Similarity Analysis (RSA; Kriegeskorte et al., 2008) to quantify neural pattern similarity during the pain empathy task. To this end, we obtained single-trial estimates by modeling trials as separate regressors (Mumford et al., 2012), time-locked to the onset of each trial’s anticipation cue (**Figure 1AB**). Events were estimated as a boxcar function with the duration set until the offset of the delivery screen (mean = 6.5 s, range = 5-8 s) and were convolved with a canonical hemodynamic response function. Rating periods (6 s) were combined within a task regressor of no interest and the six realignment parameters were appended to capture the effects of head motion. A high-pass filter with a cutoff at 128 s was applied. This resulted in 60 single-trial beta estimates per subject that were used for subsequent RSA.

Second, we used Psychophysiological Interaction analysis (PPI; Friston et al., 1997) to test connectivity during the pain empathy task. We thus adapted the first-level analysis such that trials of each condition were collapsed into four task regressors of interest (i.e., self pain, self no pain, other pain, other no pain; Rütgen et al., 2015b). The remaining regressors were modeled identically to the analysis above and contrasts were computed to assess connectivity differences between pain and no pain conditions (i.e., self pain > self no pain, other pain > other no pain).

### Representational Similarity Analysis (RSA)

We moved a spherical searchlight (Kriegeskorte et al., 2006; see also Wagner et al., 2016) with a radius of 8 mm (251 voxels) throughout the brain volume while only considering searchlights that contained at least 30 gray matter voxels. Single-trial beta estimates from voxels within a given searchlight were extracted and reshaped into a trial × voxel matrix, whereby trials were sorted according to the four experimental conditions (i.e., self pain, self no pain, other pain, other no pain). Data were *z*-scored across trials to remove mean activation differences, and voxel patterns of each trial were correlated with the voxel patterns of all other trials, resulting in a trial × trial similarity matrix. This matrix was then Fisher’s *z*-transformed and pattern similarity scores were calculated by averaging across the respective quadrants of the similarity matrix. First, we assessed pattern similarity for painful and non-painful electrical stimulation, separately for self-vs. other-directed conditions ([self pain × self no pain], [other pain × other no pain]; **Figure 1B**). Second, and central to our hypothesis, we assessed pattern similarity between self- and other-directed trials during painful vs. non-painful electrical stimulation ([self pain × other pain], [self no pain × other no pain]; **Figure 1C**). The pattern similarity values were then assigned to each searchlight’s center voxel, yielding four 3-dimensional whole-brain pattern similarity maps per subject.

Group-level significance was tested with paired-sample *t*-tests, comparing pattern similarity (1) of self-vs. other-directed trials during painful and non-painful electrical stimulation (i.e., [self pain × self no pain] vs. [other pain × other no pain]), and (2) between self- and other-directed trials during painful vs. non-painful electrical stimulation (i.e., [self pain × other pain] vs. [self no pain × other no pain]). We applied cluster-inference with a cluster-defining threshold of *p* < 0.001 and a cluster-probability threshold of *p* < 0.05 FWE-corrected for multiple comparisons for all analyses. The corrected cluster size threshold (i.e., the spatial extent of a cluster that is required in order to be labeled as significant) was calculated using the SPM extension “CorrClusTh.m” and the Newton-Raphson search method (script provided by Thomas Nichols, University of Warwick, United Kingdom, and Marko Wilke, University of Tübingen, Germany; http://www2.warwick.ac.uk/fac/sci/statistics/staff/academic-research/nichols/scripts/spm/). Anatomical nomenclature was obtained from the Laboratory for Neuro Imaging (LONI) Brain Atlas (LBPA40, http://www.loni.usc.edu/atlases/; Shattuck et al., 2008).

#### Association of pattern similarity with confederate similarity and trait empathy

We further tested if pattern similarity between self- and other-directed trials during the pain empathy task was associated with the perceived similarity of the confederate, as well as with the different aspects of trait empathy (i.e., IRI subscales). We calculated individual difference maps, subtracting pattern similarity between self- and other-related non-painful trials from pattern similarity between self- and other-related painful trials in a voxel-wise manner (i.e., [self pain × other pain] minus [self no pain × other no pain]). These pattern similarity-difference maps were then submitted to separate linear regression analyses with individual ratings of perceived confederate similarity or trait empathy added as a covariate of interest.

### Connectivity analysis

We performed two PPI analyses (contrasts self pain > self no pain, other pain > other no pain) with a seed placed within the anatomical boundaries of the left hippocampus (based on the Automatic Anatomical Labeling atlas; Tzourio-Mazoyer et al., 2002). The first eigenvector of the seed’s time course was extracted (i.e., the physiological factor) and adjusted for average activation during the task using an *F*-contrast. The eigenvector was then convolved with the respective task condition (i.e., the psychological factor), and connectivity positively related to this interaction was investigated. Contrasts were then submitted to one-sample *t*-test for random-effects, second-level analysis.

## Results

### Pattern similarity during self- and other-directed electrical stimulation

As the first analysis step, we investigated general differences between the neural representations of self-compared to other-directed painful and non-painful electrical stimulation across trials using whole-brain multivoxel pattern similarity (i.e., the main effect of stimulation target; [self pain × self no pain] vs. [other pain × other no pain]). Results revealed increased pattern similarity within bilateral insula, dACC/aMCC, right primary motor and somatosensory cortices (note that electrical stimulation was delivered to the left hand), and occipital regions during self-compared to other-directed electrical stimulation (contrast [self pain × self no pain] > [other pain × other no pain]; **Figure 2A, Table 1**). Conversely, during other-compared to self-directed electrical stimulation, we found increased pattern similarity in bilateral fusiform gyrus and surrounding inferior temporal cortex, hippocampus and parahippocampal gyrus, striatum and subgenual ACC, as well as anterior and posterior midline structures (contrast [other pain × other no pain] > [self pain × self no pain]; **Figure 2B, Table 1**).

**Figure 2.**
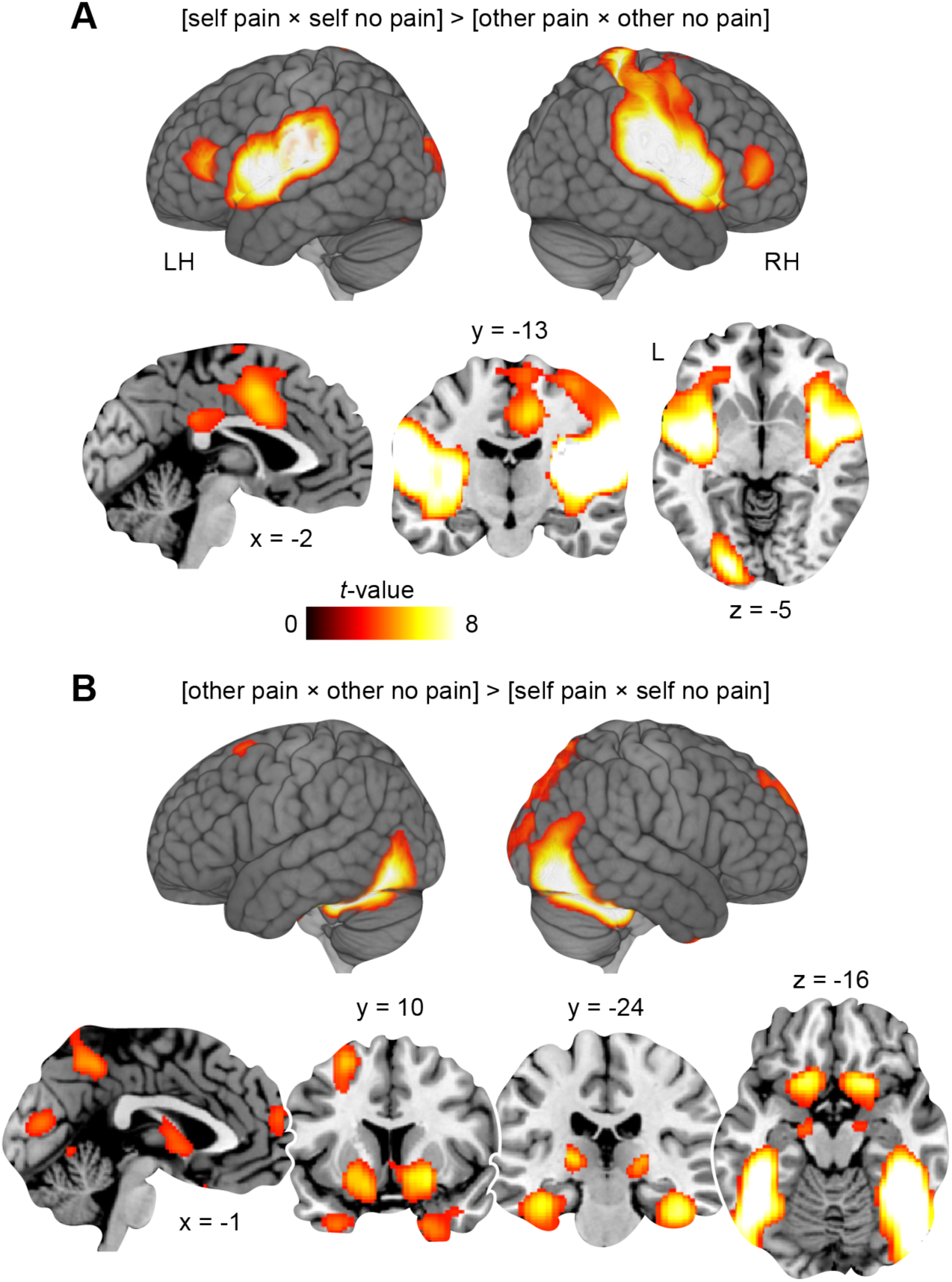
Pattern similarity during self- and other-directed electrical stimulation. Increased pattern similarity during **(A)** [self pain × self no pain] > [other pain × other no pain] and **(B)** [other pain × other no pain] > [self pain × self no pain]. Results are shown at *p* < 0.001 (*p* < 0.05, FWE-corrected at cluster-level; see also **Table 1**). L, left; LH, left hemisphere; RH, right hemisphere.

**Table 1.**
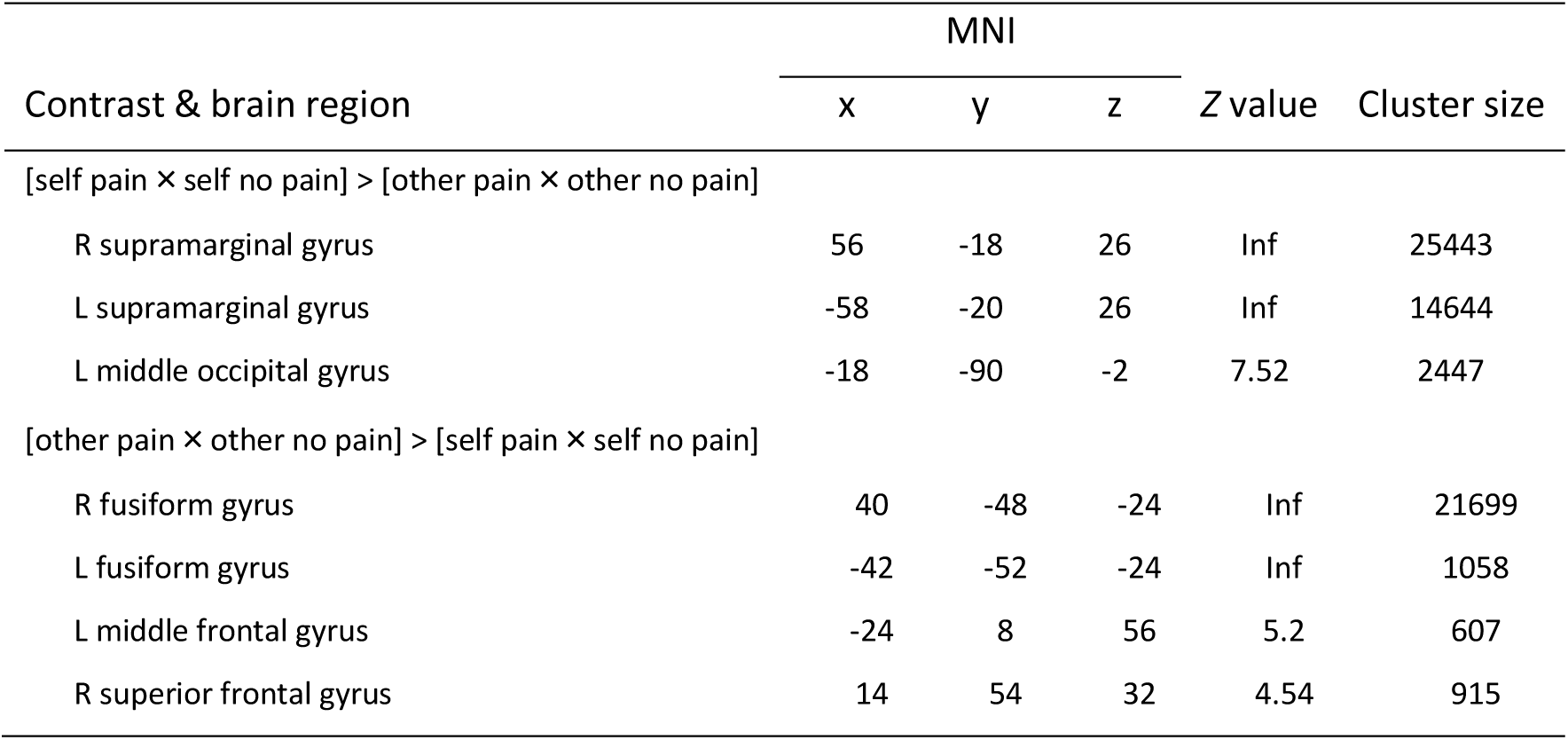
Pattern similarity during self- and other-directed electrical stimulation. MNI coordinates represent the location of peak voxels. We report the first local maximum within each cluster. Effects were tested for significance using cluster inference with a cluster-defining threshold of *p* < 0.001 and a cluster probability of *p* < 0.05, FWE-corrected for multiple comparisons (critical cluster size: 605 voxels). L, left; R, right.

### Pattern similarity of first-hand pain and pain empathy

The next step comprised the critical test of our main hypothesis, i.e., if mental simulation contributes to pain empathy, participants should utilize first-hand pain experiences to evaluate the pain of another individual. This should be associated with increased multivoxel pattern similarity (a proxy for similar neural representations) in the hippocampus and surrounding MTL. Furthermore, we expected increased pattern similarity in distributed regions known to play a role in mental simulation and pain empathy such as the MPFC, PCC, TPJ, dACC/aMCC, and the anterior insula. We reasoned that empathic processing should be increased during pain and thus tested our predictions by contrasting the pattern similarity of self- and other-directed electrical stimulation between painful and non-painful trials (i.e., [self pain × other pain] vs. [self no pain × other no pain]).

Results showed increased pattern similarity between self- and other-directed painful compared to non-painful electrical stimulation within the left hippocampus, bilateral retrosplenial cortex, extending into the fusiform gyrus and inferior temporal cortex, occipital regions, bilateral TPJ, bilateral anterior insula and the right primary motor cortex (contrast [self pain × other pain] > [self no pain × other no pain]; **Figure 3A, Table 2**). Effects for self-other similarity during non-painful relative to painful electrical stimulation were located in left visual and somatosensory cortices (contrast [self no pain × other no pain] > [self pain × other pain]; not shown in figure, see **Table 2**). Thus, as expected, the multivoxel patterns between first-hand pain and pain empathy appeared similar in the hippocampus, inferior temporal and retrosplenial cortex, TPJ, primary motor cortex and anterior insula.

**Figure 3.**
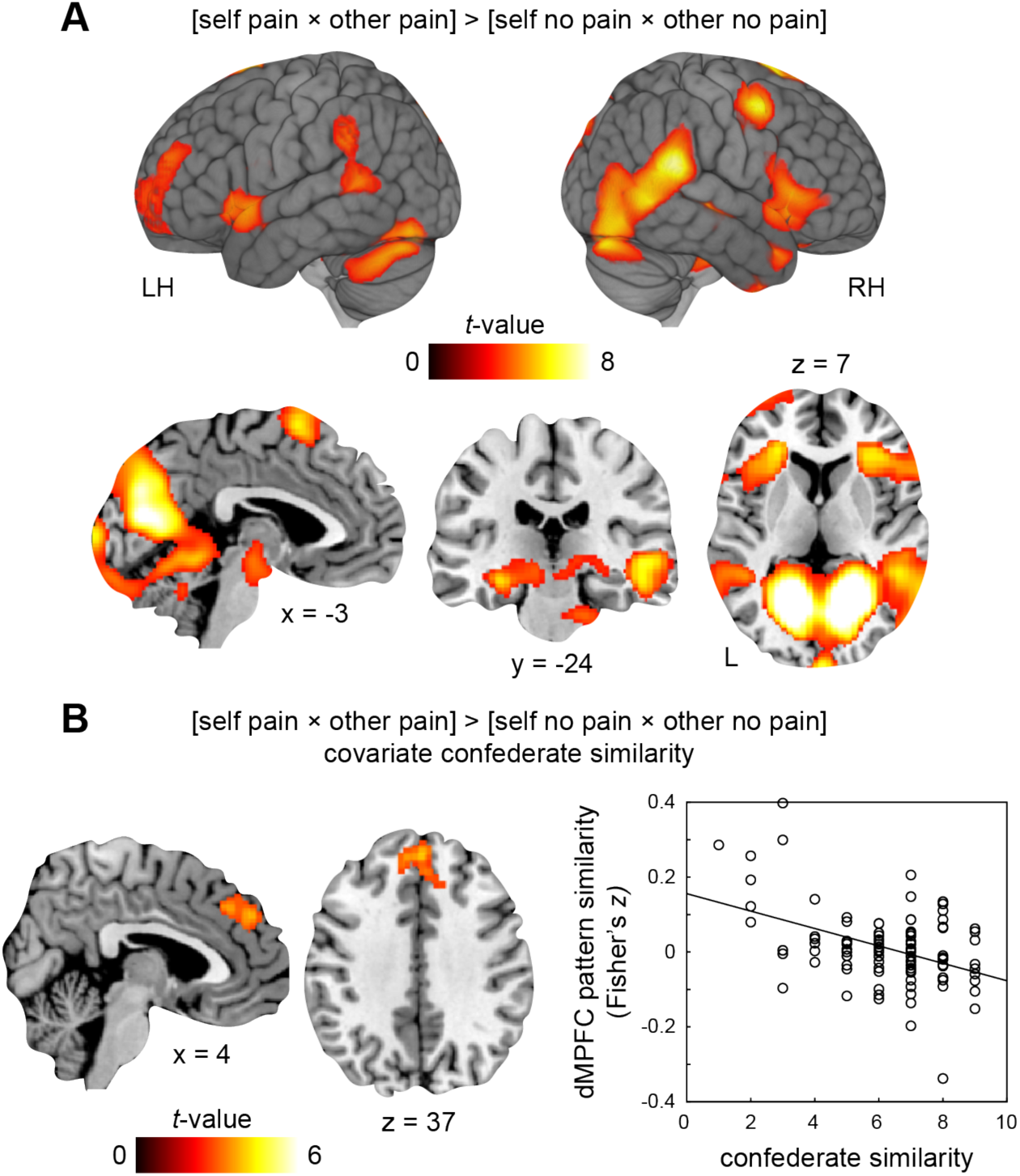
Pattern similarity of first-hand pain and pain empathy, and relation with trait empathy. **(A)** Increased pattern similarity during [self pain × other pain] > [self no pain × other no pain]. **(B)** Increased pattern similarity between first-hand pain and pain empathy was coupled to lower perceived confederate similarity across participants. For visualization purposes, the scatter plot shows the relationship between individual ratings of perceived confederate similarity and pattern similarity (Fisher’s *z* values), extracted from the significant cluster within the dMPFC. Results for both **(A)** and **(B)** are shown at *p* < 0.001 (*p* < 0.05, FWE-corrected at cluster-level; see also **Table 2**). L, left; LH, left hemisphere; RH, right hemisphere.

**Table 2.** Pattern similarity of first-hand pain and pain empathy, and relation with trait empathy. MNI coordinates represent the location of peak voxels. We report the first local maximum within each cluster. Effects were tested for significance using cluster inference with a cluster-defining threshold of *p* < 0.001 and a cluster probability of *p* < 0.05, FWE-corrected for multiple comparisons (critical cluster sizes: paired-samples *t*-test, 591 voxels; linear regression, 196 voxels). L, left; R, right.

| Contrast & brain region | MNI |  |  | Z value | Cluster size |
| --- | --- | --- | --- | --- | --- |
|  | x | y | z |  |  |
| [self pain × other pain] > [self no pain × other no pain] |  |  |  |  |  |
| L lingual gyrus | -14 | -66 | 8 | Inf | 34625 |
| R inferior frontal gyrus | 38 | 30 | 2 | 6.1 | 4508 |
| R superior frontal gyrus | 12 | 6 | 66 | 6.03 | 2068 |
| L middle frontal gyrus | -26 | 32 | 6 | 5.48 | 2719 |
| L middle temporal gyrus | -48 | -46 | 6 | 4.49 | 963 |
| L middle frontal gyrus | -40 | 54 | 24 | 4.37 | 1036 |
| [self no pain × other no pain] > [self pain × other pain] |  |  |  |  |  |
| L middle occipital gyrus | -22 | -102 | -6 | 6.22 | 891 |
| L postcentral gyrus | -36 | -26 | 56 | 5.64 | 2181 |
| [self pain × other pain] > [self no pain × other no pain],<br>covariate confederate similarity, negative effect |  |  |  |  |  |
| R superior frontal gyrus | 2 | 46 | 36 | 4.52 | 553 |

### Pattern similarity and relation to perceived confederate similarity

It has been repeatedly posited and partly confirmed that perceived similarity between self and other should be conducive to higher empathy and affective simulation (see for example Majdandžić et al., 2016). Thus, we explored whether pattern similarity of first-hand pain and pain empathy might scale with how similar participants perceived the confederate. We tested this by assessing the linear cross-participant relationship between whole-brain pattern similarity of self- and other-directed painful (compared to non-painful) stimulation with confederate similarity which was rated post-experimentally. We found that increased pattern similarity was indeed negatively associated with confederate similarity in the dorsal MPFC (**Figure 3B, Table 2**). In other words, more similar multivoxel patterns in the dorsal MPFC during first-hand pain and pain empathy were correlated with lower perceived confederate similarity across participants. No other brain region showed a significant negative or positive relationship between pattern and confederate similarities. Furthermore, there was no significant association between pattern similarity and aspects of trait empathy (i.e., subscales of the IRI). To conclude, pattern similarity between first-hand pain and pain empathy was modulated by perceived confederate similarity within the dorsal MPFC.

### Hippocampal-neocortical connectivity during first-hand pain and pain empathy

We next reasoned that if participants used representations of their previous first-hand pain experiences to evaluate the pain of others, then this should be paralleled by the recall of recent information from memory. On a neural level, this should be indexed by increased hippocampal coupling with neocortical regions that were also involved when experiencing pain first-hand. Above, we reported increased pattern similarity between first-hand pain and pain empathy within the hippocampus, whereby results appeared left lateralized (**Figure 3A, Table 2**). We thus placed a seed within the anatomical boundaries of the left hippocampus in order to test connectivity during self- and other-directed painful compared to non-painful electrical stimulation.

First, we found increased hippocampal coupling with bilateral insula, dACC/aMCC, thalamus, right primary motor and somatosensory cortices, lateral prefrontal and occipital regions when participants received painful electrical stimulation themselves (contrast self pain > self no pain; **Figure 4A, Table 3**). Second, results showed increased hippocampal connectivity with left fusiform gyrus and right primary motor cortex when painful electrical shocks were delivered to the confederate (contrast other pain > other no pain; **Figure 4B, Table 3**). Thus, hippocampal-neocortical coupling during pain empathy engaged regions that were also involved when experiencing pain first-hand.

**Figure 4.**
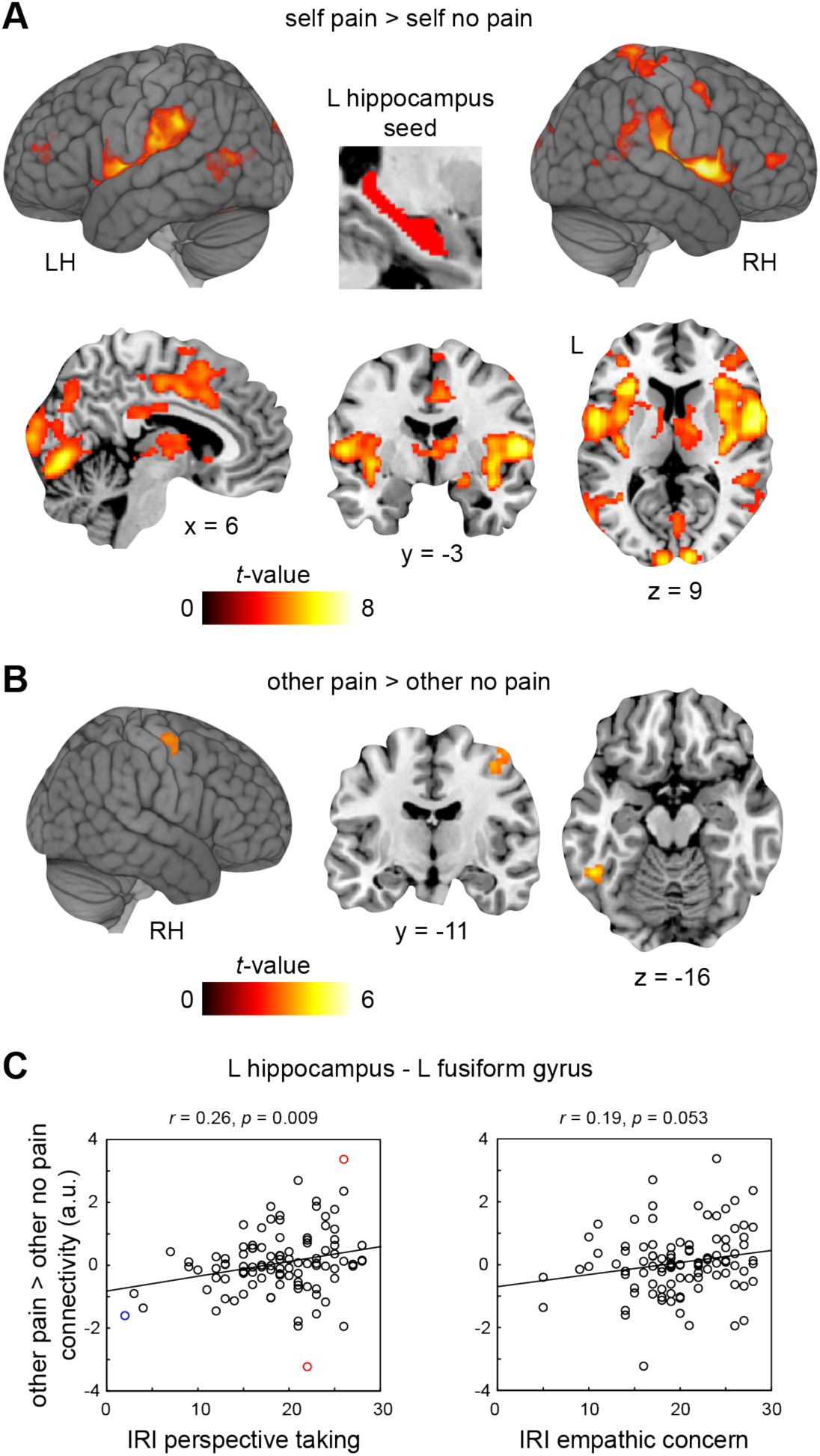
Hippocampal connectivity and association with trait empathy. Hippocampal connectivity increases during **(A)** self pain > self no pain, and **(B)** other pain > other no pain. The anatomical seed region within the left hippocampus is marked in red. Results are shown at *p* < 0.001 (*p* < 0.05, FWE-corrected at cluster-level; see also **Table 3**). **(C)** *Left panel:* Connectivity of the left hippocampus with the left fusiform gyrus (a.u., arbitrary units) showed a positive relationship with perspective taking (i.e., IRI subscale). Thus, stronger coupling was associated with higher scores in perspective taking across subjects. The correlation remained robust when removing three outliers (mean ± 3 standard deviations; one outlier in IRI perspective taking, two outliers in connectivity; *r* = 0.22, *p* = 0.026, bootstrapped 95% confidence interval based on 5000 samples [0.053 0.4]). *Right panel:* There was a trend towards increased coupling at higher levels of empathic concern across participants. L, left; LH, left hemisphere; RH, right hemisphere.

**Table 3.** Hippocampal connectivity and association with trait empathy. MNI coordinates represent the location of peak voxels. We report the first local maximum within each cluster. Effects were tested for significance using cluster inference with a cluster-defining threshold of *p* < 0.001 and a cluster probability of *p* < 0.05, FWE-corrected for multiple comparisons (critical cluster sizes: self pain > self no pain, 96 voxels; other pain > other no pain, 82 voxels). L, left; R, right.

| Contrast & brain region | MNI |  |  | Z value | Cluster size |
| --- | --- | --- | --- | --- | --- |
|  | x | y | z |  |  |
| self pain > self no pain |  |  |  |  |  |
| R precentral gyrus | 56 | 8 | 4 | 6.94 | 13816 |
| R middle occipital gyrus | 12 | -98 | 10 | 6.46 | 3803 |
| L cingulate gyrus | -2 | 10 | 36 | 5.31 | 2186 |
| R superior parietal gyrus | 24 | -40 | 76 | 5.18 | 854 |
| R lingual gyrus | 24 | -68 | -4 | 5.12 | 213 |
| L middle temporal gyrus | -58 | -64 | 12 | 4.86 | 442 |
| L middle frontal gyrus | -36 | 42 | 10 | 4.6 | 194 |
| R precentral gyrus | 52 | 0 | 54 | 4.53 | 272 |
| Cerebellum | -26 | -62 | -22 | 4.47 | 288 |
| R inferior frontal gyrus | 40 | 42 | 8 | 4.17 | 196 |
| R lingual gyrus | 22 | -54 | 0 | 4.16 | 154 |
| L angular gyrus | -36 | -52 | 38 | 3.94 | 102 |
| other pain > other no pain |  |  |  |  |  |
| L inferior temporal gyrus | -48 | -50 | -16 | 4.25 | 83 |
| R precentral gyrus | 44 | -6 | 62 | 3.78 | 137 |

### Hippocampal-neocortical connectivity and association with trait empathy

Last, we examined if hippocampal-neocortical coupling scaled with confederate similarity or aspects of trait empathy. Connectivity between the hippocampus and the left fusiform gyrus positively correlated with individual differences in perspective taking (i.e., the IRI subscale). Put differently, participants who scored higher on perspective taking showed stronger hippocampal-fusiform connectivity during pain empathy (*r* = 0.26, *p* = 0.009, bootstrapped 95% CI based on 5000 samples [0.08 0.44]; **Figure 4C**, left panel). Furthermore, this connectivity profile showed a trend towards increased coupling at higher levels of empathic concern across participants (i.e., IRI subscale, *r* = 0.19, *p* = 0.053; **Figure 4C**, right panel). There was no significant association of empathy-related hippocampus-fusiform gyrus connectivity with confederate similarity (*p* = 0.557), and no significant association with the remaining IRI subscales (IRI personal distress: *p* = 0.701, IRI fantasy: *p* = 0.098). In addition, there was no significant relationship between hippocampus-primary motor cortex connectivity during pain empathy and any of the behavioral measures (confederate similarity: *p* = 0.976, IRI personal distress: *p* = 0.697, IRI perspective taking: *p* = 0.313, IRI empathic concern: *p* = 0.950, IRI fantasy: 0.946). In summary, increases of hippocampal connectivity with the fusiform gyrus during pain empathy were larger in participants with higher self-reported perspective taking skills, and empathic concern (by trend).

## Discussion

The goal of the present study was to investigate potential contributions of hippocampal-neocortical representations and connectivity to pain empathy, potentially reflecting mental simulation processes. Our analysis approach revealed four main findings: First, we found increased pattern similarity (i.e., shared neural representations) between first-hand pain and pain empathy within the hippocampus and a wider neocortical network, including inferior temporal and retrosplenial cortex, TPJ, primary motor cortex and anterior insula. Second and partly replicating prior work (Mitchell et al., 2006; Tamir and Mitchell, 2010), we showed that increased pattern similarity between first-hand pain and pain empathy within the dorsal MPFC was coupled to lower perceived confederate similarity across participants. Third, results demonstrated that hippocampal-neocortical coupling during pain empathy engaged regions that were also involved when experiencing pain first-hand. Fourth, hippocampal connectivity with the fusiform gyrus during pain empathy was larger at higher self-reported skills in perspective taking. These findings suggest that mental simulation might contribute to pain empathy via hippocampal-neocortical pattern similarity and connectivity, partially affected by personality traits and the similarity of the observed individual.

Our main hypothesis was that if mental simulation processes support pain empathy, participants should utilize first-hand pain experiences as a model to evaluate the pain of another individual. This should be based on similar neural representations, approximated by multivoxel pattern similarity between first-hand pain and pain empathy. Specifically, we expected increased pattern similarity within the hippocampus and surrounding MTL structures, as well as within a wider neocortical set of regions important for mental simulation (i.e., including MPFC, PCC, and TPJ; Frith and Frith, 1999; Amodio and Frith, 2006; Uddin et al., 2007) and pain empathy (i.e., including dACC/aMCC and anterior insula; Lamm et al., 2011). Results indeed showed increased pattern similarity within the hippocampus, extending into inferior temporal and retrosplenial cortices, TPJ, primary motor cortex and anterior insula (**Figure 3A**). The hippocampus is regarded as key player for successfully remembering past (Scoville and Milner, 1957; Rugg and Vilberg, 2012; Kim, 2016) and simulating future (Addis et al., 2007; Szpunar et al., 2014; McCormick et al., 2018) or fictitious events (Hassabis et al., 2007a), and it therefore seems vital for mental construction (Schacter and Addis, 2007; Hassabis and Maguire, 2009; Bird et al., 2010; Summerfield et al., 2010) and self-projection (Buckner and Carroll, 2007; Kurczek et al., 2015). Hippocampal lesions in human patients, for instance, were shown to impair the ability to imagine new experiences (Klein et al., 2002; Hassabis et al., 2007b), whereby specific deficits may differ depending on the lesion type and its exact location (Addis and Schacter, 2011). Importantly, Beadle and colleagues (2013) reported lower trait empathy in patients with focal hippocampal damage, along with no effect of empathy induction on empathy ratings or prosocial behavior (Beadle et al., 2013; and see also Rushby et al., 2016). These findings collectively suggest a role of the hippocampus in empathy and mental simulation. Moreover, the MPFC (Benoit et al., 2010, 2014; Kurczek et al., 2015; Schurz et al., 2015; Bertossi et al., 2016b, 2016a; Barry et al., 2019) and PCC (including precuneus and retrosplenial cortex; Vann et al., 2009; Summerfield et al., 2010; Irish et al., 2015; Schurz et al., 2015; Ramanan et al., 2018) are considered pivotal for mental construction and self-projection as well, and are also engaged during theory of mind (Frith and Frith, 1999, 2006; Uddin et al., 2007; Mar, 2011). The latter additionally involves the TPJ, possibly signaling perspective taking to simulate the mental stance of another person (Saxe and Kanwisher, 2003; Saxe and Wexler, 2005; Saxe and Powell, 2006). To summarize, the hippocampus, MPFC, PCC, and TPJ constitute a distributed set of brain regions associated with memory, mental construction and simulation, self-projection, and theory of mind. Here, we observed that these regions support pain empathy as well, holding similar representations when experiencing pain first-hand and when observing pain in another individual.

Empathy, however, goes beyond mere mental simulation as it also incorporates affect sharing and self-other distinction (Lamm et al., 2019). While the latter might be achieved by neural processing within the TPJ (Saxe and Kanwisher, 2003), affect sharing may depend on shared representations during the first-hand (pain) experience and empathy for it (Bastiaansen et al., 2009; Singer and Lamm, 2009; Decety, 2010). Previously, this has been linked to neural processing within the dACC/aMCC and anterior insula (Corradi-Dell’Acqua et al., 2011, 2016; Lamm et al., 2011; Rütgen et al., 2018; Carrillo et al., 2019), including an initial, univariate analysis of the current study (Rütgen et al., 2015b). Using a novel multivariate analysis approach, we here partly confirmed and extended these previous findings: First, we found increased pattern similarity within the dACC/aMCC and insula during first-hand pain (compared to pain empathy), while pain empathy (compared to first-hand pain) was associated with regions such as hippocampus, inferior temporal cortex, including the fusiform gyrus, MPFC, and PCC (**Figure 2AB**). Second, we tackled the question of shared neural representations between first-hand pain and pain empathy and found increased pattern similarity in a set of regions, including bilateral anterior insula (**Figure 3A**). The insula was associated with pain processing (Downar et al., 2003; Legrain et al., 2011) and interoceptive awareness (Craig, 2002, 2009), and similar neural representations might thus signal affective sharing with the observed individual (Singer et al., 2004; Lamm et al., 2011). Somewhat surprising and unexpectedly, we could not identify increased pattern similarity within the dACC/aMCC. While previous work showed overlapping neural assemblies representing first-hand pain as well as pain empathy within the rodent dACC/aMCC (although lacking a direct or closely matched comparison between painful and non-painful events; Sakaguchi et al., 2018; Carrillo et al., 2019), others reported a domain-general role of the dACC/aMCC in empathic processing of photographs of human body parts during painful and non-painful situations (Gu et al., 2010). Also, anterior insula engagement in empathy has been linked to emotional (self-)awareness and interoception of the associated affective states, whereas dACC/aMCC seems to be recruited by physiological regulatory processes, including homeostasis, rather than representing the feelings of the self or the other person (Craig, 2009; Lamm et al., 2011). We thus speculate that the dACC/aMCC might reflect neural processes related to self and other during electrical stimulation independent of stimulation intensity, and suggest future studies to clarify the exact role of the human dACC/aMCC in pain empathy. To conclude, we found shared neural representations between first-hand pain and pain empathy in the anterior insula that, together with hippocampus, MPFC, PCC, and TPJ, appear to support empathy, potentially via affect sharing and mental simulation.

Empathic processing, and associated mental simulation, might depend on how similar the individual is perceived to oneself. Our results demonstrate that lower perceived confederate similarity was coupled to increased pattern similarity between first-hand pain and pain empathy in the dorsal MPFC (**Figure 3B**). The dorsal MPFC was previously implicated in self-projection and perspective taking (Kurczek et al., 2015; Schurz et al., 2015), self-inhibition/other-enhancement during mental simulation (Majdandžić et al., 2016), and prosocial behavior (Waytz et al., 2012). Mitchell and colleagues (2006) showed that judgements of similar and dissimilar others were associated with activation changes in ventral and dorsal MPFC, respectively (Mitchell et al., 2006). To illustrate this further, Tamir and Mitchell (2010) demonstrated that MPFC activation was greater with increasing discrepancy between self- and other-related judgements (Tamir and Mitchell, 2010), and multivariate patterns within a similar region were shown to scale with how fine-grained or distinct self-compared to other-related mental states were described (Thornton et al., 2019). In line with these findings, our analyses also suggest a specific role of the dorsal MPFC in mental simulation, in particular if the other individual is perceived as less similar to oneself. In this context, it should also be noted that we did not find a significant modulation of pattern similarity between first-hand pain and pain empathy by confederate similarity in the hippocampus (but see Perry et al., 2011), PCC, or TPJ. In light of the inherent challenges of interpreting null findings, it thus remains an open question of whether and how these areas are relevant for the simulation of persons we perceive as more similar to us (see also Majdandžić et al., 2016 for discussion of related challenges in demonstrating the complex relationship between self-other similarity, empathy, and prosocial behavior). Next, we hypothesized that if mental simulation supports pain empathy, then this should be dovetailed with memory recall of the recently experienced painful stimulation. We found increased hippocampal connectivity with left fusiform gyrus and right primary motor cortex during pain empathy (**Figure 4B**). Indeed, the right primary motor cortex was also engaged when participants experienced electrical stimulation to the left hand (**Figure 2A**), and this region also showed increased pattern similarity between first-hand pain and pain empathy (**Figure 3A**). Memories are assumed to be stored in distributed neocortical networks (Marr, 1970; Frankland and Bontempi, 2005). The hippocampus is thought to coordinate (recent) memory retrieval through coupling with neocortical regions that were engaged during the actual experience (Frankland and Bontempi, 2005; Takashima et al., 2009). Increased hippocampal connectivity with the right primary motor cortex during pain empathy might thus be related to the recall of recently experienced pain. Furthermore, participants with higher trait measures in perspective taking showed stronger hippocampal coupling with the fusiform gyrus during pain empathy. We speculate that the fusiform gyrus contributed to simulation processes, presumably coding for the content of visual imagination (O’Craven and Kanwisher, 2000; Pearson, 2019). Note though that this constitutes a reverse inference, in need of more direct future validation by studies incorporating measures of, for example, individual differences in self-reported visual imagery skills.

Empathy, including affect sharing (Hein et al., 2010) or mental simulation (Waytz et al., 2012; Gaesser and Schacter, 2014; Gaesser et al., 2015, 2017, 2018, 2019), might ultimately motivate prosocial behavior. Gaesser and colleagues (2014), for example, showed that episodic simulation and memory of helping another individual in need positively affected the willingness to help others (Gaesser and Schacter, 2014; Gaesser et al., 2015). Prosocial behavior appears increased the more vividly participants engage in mental simulation or memory recall of helping behavior (Gaesser et al., 2017, 2018, 2019), and this involved the MTL and TPJ (Gaesser et al., 2019). While the link between empathy, mental simulation, and prosocial behavior warrants further investigation, results suggest that mental simulation might contribute to empathy through neural processes in hippocampal-neocortical ensembles.

Last but not least, several limitations of our approach need to be addressed. First, our approach draws on an indirect investigation of mental simulation and shared representations to pain empathy by capitalizing on hippocampal-neocortical contributions. Our results confirm our *a priori* expectations of increased hippocampal-neocortical pattern similarity and connectivity. Although an interpretation of the results in terms of mental simulation processes and memory recall during pain empathy appears likely (Frith and Frith, 1999; Buckner and Carroll, 2007; Hassabis and Maguire, 2009; Gaesser et al., 2019; Lamm et al., 2019), conclusions should be drawn with caution. Future studies should investigate mental simulation and empathy within the same study and could then directly link both. Nevertheless, our results provide novel insights in hippocampal processing during empathy, corroborating previous findings on empathy deficits in amnesia (Beadle et al., 2013) and traumatic brain injury (Rushby et al., 2016). Second, we tested the relationship between shared representations (approximated by representational pattern similarity between first-hand pain and pain empathy) and confederate similarity. We did not manipulate confederate similarity but instead assessed it post-experimentally. Future studies may systematically manipulate confederate similarity for a more detailed investigation.

To conclude, we investigated hippocampal-neocortical contributions to pain empathy and linked them to mental simulation processes. We found increased pattern similarity between first-hand pain and pain empathy within the hippocampus and a set of neocortical regions, including the inferior temporal and retrosplenial cortex, TPJ, anterior insula, and primary motor cortex. While neural representations in these regions were unaffected by confederate similarity, pattern similarity in the dorsal MPFC was increased the more dissimilar the other individual was perceived. During pain empathy, hippocampal coupling with the primary motor cortex and fusiform gyrus was increased, whereby hippocampal-fusiform connectivity was stronger at higher individual perspective taking skills. Together, these findings highlight the contributions of hippocampal-neocortical representations and connectivity to pain empathy, partially affected by personality traits and the similarity of the other individual in pain. This might potentially indicate mental simulation and memory recall during empathy, achieved through shared neural representations and hippocampal-neocortical interactions, and could have important practical implications for empathy in patients suffering from hippocampal damage or fronto-temporal dementia.

## Acknowledgements

The authors would like to thank Allan Hummer and Christian Windischberger from the Medical University of Vienna, as well as Daniel Graf, Bernadette Hippmann, Julia Hebestreit, Alexander Kudrna, and Andreas Martin from the University of Vienna for assistance with functional MRI measurements; Fritz Zimprich (Medical University of Vienna) for medical support; and Andreas Gartus (University of Vienna) for technical support. The study was supported by the Vienna Science and Technology Fund (WWTF; Project CS11-016), and the Austrian Science Fund (FWF; Project P32686, awarded to CL, MR, and ICW).

